# Recurrent Escape from Osimertinib-Induced Senescence Promotes Genomic Instability Associated with Therapeutic Resistance

**DOI:** 10.64898/2026.02.07.704600

**Authors:** Nazia S Jamil, Qualia Hooker, Nadjet Cornejal, H Dean Hosgood, Hayley M McDaid

## Abstract

Acquired resistance to osimertinib remains a major challenge in treating EGFR-mutant (EGFR+) Non-Small-Cell Lung Cancer (NSCLC). Although most patients initially respond to treatment, relapses are universal, even after prolonged remission during which tumor dormancy occurs. Here, we show that osimertinib induces and maintains senescence in EGFR+ NSCLC. Importantly, osimertinib does not kill senescent cells; however, following drug withdrawal, cells escape and resume proliferation. To examine the consequences of recurrent senescence and escape on resistance, we generated four isogenic cell lines clonally expanded through sequential cycles of **Os**imertinib-**I**nduced **S**enescence (OsIS). Phylogenetic reconstruction based on *de novo* somatic variants revealed that these lines form four distinct evolutionary clades with varying degrees of osimertinib resistance. All had elevated tumor mutational burden with distinct single-nucleotide and copy-number variants, and without acquisition of tertiary *EGFR* mutations or *MET* amplification. Resistance was predominately associated with chromosomal instability characterized by extensive loss of heterozygosity, high copy-number alteration burden, and mutational signatures consistent with replication-associated DNA damage and repair. A second resistance genotype exhibited extreme focal amplifications with breakage-fusion-bridge-like genome remodeling. Despite profound genomic instability, targeting DNA repair or replication stress pathways was ineffective, whereas sensitivity to platinum-based chemotherapy was retained across clades. Collectively, these findings indicate that recurrent senescence escape drives osimertinib resistance through widespread genomic instability and is most effectively treated by cytotoxic strategies rather than pathway-targeted approaches.

**Significance:** Although most patients with EGFR+ lung cancer relapse after osimertinib therapy, only a small fraction of cases are explained by on-target resistance mutations. This study shows that recurrent cycles of osimertinib-induced senescence and escape promote resistance through chromosomal instability, identifying dormant cells as critical reservoirs for relapse.

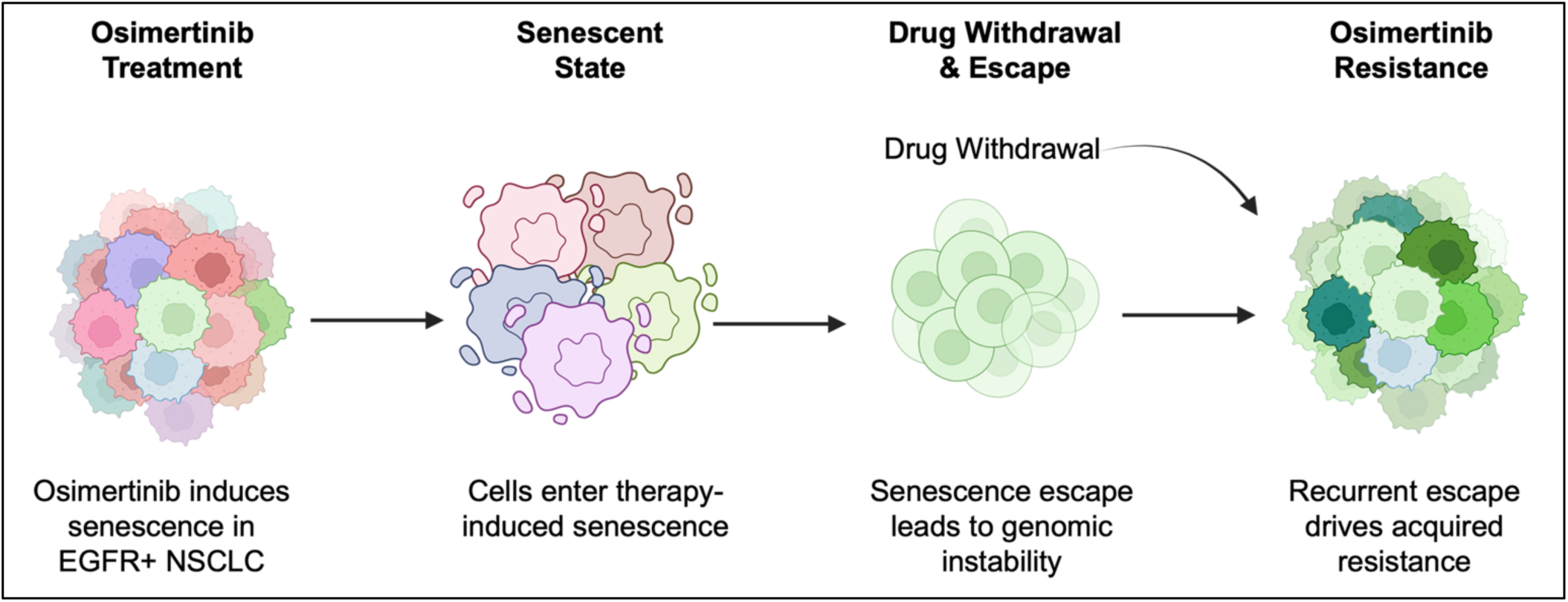

## Introduction

Osimertinib has superior efficacy relative to first- and second-generation EGFR Tyrosine Kinase Inhibitors (TKIs), establishing it as the global standard of care for first-line treatment of EGFR⁺ Lung Adenocarcinoma (LUAD) (1, 2). However, acquired resistance is essentially universal, with median progression-free survival of 16.7 months in FLAURA2 (3, 4). Despite progression, osimertinib is often continued with platinum-based chemotherapy, resulting in significantly improved outcome compared with chemotherapy alone (5, 6).

One well-documented mechanism of clinical resistance is the acquisition of tertiary mutations in *EGFR* that impair osimertinib binding. Mutation at C797X occurs in 6–8% of patients following first-line osimertinib (7, 8), with less common variants at L718Q/V, G796S/R, L792H/F, and exon 20 insertions accounting for approximately 4% of cases (7). Fourth-generation EGFR inhibitors targeting C797S are currently in development and early clinical trials are ongoing (9). Another alteration associated with clinical relapse is *MET* amplification, detected in approximately 15–20% of patients. However, these are typically subclonal mutations that co-occur with widespread copy-number changes, consistent with increased genomic instability (10). Notably, MET inhibitors are not FDA-approved for MET-amplified, EGFR+ NSCLC after progression on osimertinib, despite multiple clinical trials (11, 12). Comprehensive genomic profiling of osimertinib-resistant tumors reveals marked genetic heterogeneity in contrast to the more homogeneous clonal architecture of treatment-naïve disease (8). Yet the timing and mechanisms by which therapy promotes this heterogeneous, recalcitrant state remain poorly understood (13).

The prolonged period of remission for adjuvant osimertinib reported in the ADAURA trial indicates durable disease control while on therapy with median disease-free survival approximating 5.5 years (14). At the tumor level, this is consistent with therapy-induced dormancy and no evidence of disease status, while recurrence that coincides with resistance to osimertinib is preceded by escape from a prolonged growth-arrested state. Cellular senescence is a stable form of proliferative arrest that is a hallmark of cancer (15), and a common outcome of cancer therapy (16, 17), including EGFR inhibition (18–20). Senescent cancer cells pose a significant threat because they exist in a transitory state and can resume proliferation (15) with more aggressive tumor biology traits (21, 22).

Resistance models based on stepwise dose escalation over time frequently select for drug-binding site mutants through persister-type survival (23, 24), but these largely ignore therapy-induced senescence as an intermediate dormant state. We hypothesize that osimertinib-induced senescence functions as an evolutionary bottleneck, and that escape from this state is coupled to the emergence of resistance. Here, we show that osimertinib induces senescence in EGFR+ NSCLC cells and that, upon drug withdrawal, cells resume proliferation. We also demonstrate that recurrent senescence escape drives genomic diversification, dominated by chromosomal instability that is associated with osimertinib resistance.

## Materials and Methods

### Cell Culture and Reagents

The EGFR+ NSCLC cell line, HCC827 have an exon 19 EGFR deletion and extensive EGFR gene amplification. They were purchased from ATCC and cultured in RPMI-1640 (Gibco) supplemented with 10% fetal bovine serum (ThermoFisher) at 37°C in a humidified atmosphere with 5% CO₂. Only cells under passage 30 were used and authentication by short tandem repeat profiling, and testing for mycoplasma contamination prior to experimentation was performed every 3 months. All anti-cancer molecules were purchased from Med Chem., solubilized in 100% DMSO and stored at -20 degrees. Cisplatin was formulated in saline every 4 weeks and stored at 4 degrees. All other general chemicals were purchased from Sigma.

### Cell Proliferation Assays & Cell Line Characterization

Cell proliferation assays were performed using sulforhodamine B (SRB) to measure total protein in adherent cells (25). Cells were seeded at 3-5 × 10^4^ cells / ml into 96-well plates, and 24h later incubated with serial dilutions of test molecules or drugs, spanning 9-18 dose levels (*n*=3-6 wells per dose level). Standard cell proliferation assays were performed for 10 days without replenishing. Additional experiments also evaluated effects at 3 and 21 days. The anti-cancer effect was computed relative to vehicle-only treated control cells and sigmoidal dose-response curves generated. Alternatively, relative change in drug response was determined by calculating AUC values from dose-response plots and computing the ratio of senescence escaped versus HCC827 lines for each compound. Zero indicated no change in sensitivity whereas higher values represent decreased AUC (greater resistance). Experiments to assess long-term effects of osimertinib were reported as drug effect without normalization since control cells continued to proliferate over the duration of the experiment and eventually died.

Doubling time was determined by seeding 500 cells in 60mm^2^ wells and counting daily until confluency. Doubling time was calculated during the exponential growth phase using the formula: Td = t × ln(2) / ln(N(t)/N₀), where t is the time interval in hours and N(t)/N₀ is the fold-change in cell number between two timepoints. Mean cell area (μm²) was determined using the ECHO Revolve microscope with integrated image analysis software from subconfluent cultures. Approximately 50 cells per clade were counted and mean area ± standard deviation reported.

### Induction of Osimertinib-Induced Senescence (OsIS)

Approximately 80-100 100mm^2^ plates of 90% confluent HCC827 cells were incubated with 0.5 µmol/L osimertinib for 2 days, followed by 0.2 µmol/L osimertinib every 2 days for a further 8 days, without media change. On day 11, cells were washed three times with 1× PBS and used for experiments. This resulted in approximately 95% cell death and surviving cells became senescent.

### Senolysis and Stability of Senescence

Osimertinib-induced senescent cells were trypsinized and replated in 96-well plates at 5000 cells/well. After 24h DMSO or saline, osimertinib (0.5 µmol/L), navitoclax (10 µmol/L), cisplatin (100 µmol/L) or pemetrexed (200 µmol/L) were added to replicate wells and viability determined by SRB after 72 hours. Percent effect was determined from the proportion of surviving cells relative to vehicle controls. In addition, to monitor the effect of long term osimertinib on senescent cell viability and the stability of proliferative arrest, cells were treated weekly with 0.5 µmol/L osimertinib, media or DMSO and imaged weekly by phase-contrast microscopy (10x magnification, four fields of view). Cell count was determined as mean count ± SD per field of view.

### Generation of Senescence-Escape Clades

After osimertinib wash-out, senescent cells escaped approximately 2-4 weeks after. A colony was selected, cloned, expanded and re-dosed with 0.5 µmol/L osimertinib for 10 days to re-induce senescence. This process was repeated three times. During the third and fourth cycles, osimertinib was increased to 1.0–2.0 µmol/L to achieve ∼95% cell death and four different phylogenies selected. Cumulative osimertinib exposure during selection was approximately 40 days over 6–7 months. Following the final clonogenic expansion, cells were maintained in the absence of osimertinib.

### Wound Healing Migration and Invasion Assay

For migration, 80,000–150,000 cells per well were seeded in ImageLock 96-well plates in 10% serum overnight, and wound closure monitored over 24 hours using the Incucyte Scratch Wound software module. For invasion, 7500 cells per well were seeded into ClearView chemotaxis plates (8 µm pores) containing 500µg/mL matrigel diluted in media containing 0.5% serum. The chemotaxis insert was transferred into a pre-loaded plate containing 20% serum and placed in an Incucyte S3 and scanned using the chemotaxis module, imaging every hr for 24h. Invasion was quantified by measuring total phase object area on the bottom normalized to initial phase object area of the top layer.

### Fluorescence In Situ Hybridization (FISH)

Interphase cells were hybridized with EGFR and CEP7 probes (Empire Genomics) and counterstained with DAPI, according to the manufacturer’s protocol. Images were acquired at 100x, and ≥50 interphase nuclei per sample were scored for EGFR signal intensity relative to CEP7.

### Immunoblot Analysis

Adherent cells were collected from proliferating, quiescent, and osimertinib-treated cultures at 6 and 10 days. Cells were lysed in Tris-SDS lysis buffer, quantified, equal proteins resolved by SDS-PAGE, and transferred onto nitrocellulose. Membranes were blocked, incubated with primary antibodies overnight at 4°C and horseradish peroxidase–conjugated secondary antibodies (see Supplementary methods). Loading was confirmed by staining membranes with Ponceau S solution Protein bands were visualized using enhanced chemiluminescence (ECL) detection reagent and detected by exposure to X-ray film.

### Whole-Exome Sequencing (WES)

Library preparation and whole-exome sequencing was performed by Novogene using purified genomic DNA. Paired-end sequencing (2 × 150 bp) was performed on an Illumina NovaSeq 6000 platform by Novogene Corporation (Sacramento, CA), achieving a mean coverage depth of >98× across targeted regions.

### Phylogenetic Tree Generation

Phylogenetic relationships were inferred from germline SNP genotypes generated by Novogene. Pairwise Jaccard distances were calculated across germline SNPs using the vegan package, and a neighbor-joining tree was constructed from the distance matrix using the ape package. The resulting cladogram was visualized in Rstudio (v.4.5.0) using ggtree.

### Copy-number analysis of low pass WGS

For structural variant and copy-number analysis, low-pass whole-genome sequencing (WGS) was performed (Azenta). The mean coverage for samples was 1.0×. Paired-end reads from HCC827 parental and resistant samples were trimmed using Trim Galore (v0.6.10) (quality threshold 20) and aligned to GRCh38 using BWA-MEM (v0.7.17). BAM files were coordinate-sorted and duplicate-marked using Picard (v2.27.5). Base quality score recalibration was performed using GATK (v4.5.0.0) BaseRecalibrator and ApplyBQSR with known sites from dbSNP, Mills, and GATK resource bundles. Final BAM files were analyzed using Picard CollectWgsMetrics and samtools. Quality metrics were aggregated using MultiQC (v1.14).

### AmpliconSuite

Analysis-ready BAM files were downsampled to 10% of reads. Detection of focal amplifications was performed using AmpliconSuite-pipeline (v1.5.0), including Amplicon Architect for reconstruction and Amplicon Classifier for structural categorization against the GRCh38 reference genome. Candidate seed regions were restricted to intervals with copy-number greater than 5 and length greater than 100,000 bp. For samples with reduced properly paired rates, --insert_sdevs 9.0 was applied to filter artifacts. Amplicons were classified as breakage-fusion-bridge (BFB), complex non-cyclic, linear or unknown. Copy number amplifications were determined from Amplicon Classifier output and copy number of specific genes plotted.

### Mutational Signatures and Allele-Specific CNV analysis in Whole-exome sequencing

Germline and somatic SNVs and indels called by Novogene were used to calculate the contribution of COSMIC SBS96 and ID mutational signatures using SigProfilerAssignment (26). Loss of heterozygosity (LOH) was inferred using FACETS (Fraction and Allele Specific Copy-Number Estimates from Tumor Sequencing) (v0.6.2) (27). SNP-pileups were generated from resistant clone BAMs normalized to parental HCC827 BAM using common germline variants (1000 Genomes phase 1). FACETS-derived CNV profiles were used as input into COSMIC SigProfilerAssignment to characterize mutational signatures associated with copy-number alterations. Figures were generated in Rstudio (v4.5.0) using ggplot2.

To obtain global copy-number profiles from whole-exome sequencing samples a normal reference was generated using UCSC hg38, Agilent SureSelect Human All Exon V6 target regions (padded), and the access-10kb.hg38.bed accessible regions file provided by CNVkit. HCC827 parental and resistant BAMs were processed through cnvkit.py batch with germline SNP calls to infer B-allele frequency and LOH. Samples were annotated as female. Copy-number segments were identified using Circular Binary Segmentation (CBS) and called using custom thresholds (log2 ratios: -1.1, -0.4, 0.3, 0.7 for loss/neutral/gain/amplification) with median centering and filtering of low-coverage regions (--drop-low-coverage). Copy-number profiles across all samples were visualized using CNVkit’s heatmap function.

### Statistical Analysis

All statistical analyses were performed using GraphPad Prism version 10.6.1 (GraphPad Software Inc.). For comparison of two groups, unpaired two-tailed Student’s *t*-tests were used. For multiple group comparisons, one-way or two-way analysis of variance (ANOVA) was performed as appropriate for the experimental design. Post-hoc tests were selected based on the comparison structure: Dunnett’s test was used when comparing multiple treatment groups to a single control group (parental HCC827), and Tukey’s test was used for all pairwise comparisons between groups. Dose-response curves were analyzed using nonlinear regression (log[inhibitor] vs. response to determine maximum effect (Emax) and area under the curve (AUC). Statistical significance was defined as *P* < 0.05, with significance levels denoted as: \**P* < 0.05, \*\**P* < 0.01, \*\*\**P* < 0.001, \*\*\*\**P* < 0.0001; ns, not significant. All data are presented as mean ± standard deviation. All experiments were performed with at least three biological replicates unless otherwise stated.

### Data Availability

WES and WGS datasets generated in this study, together with analytic code are available from the corresponding author upon request.

## Results

### Osimertinib Induces and Maintains Senescence in EGFR Mutant NSCLC

To investigate the effect of a single dose of osimertinib on cell fate, HCC827 cells harboring an exon 19 deletion in EGFR, were treated with a range of concentrations and cell viability assessed at 3, 10, and 21 days without media replacement. Osimertinib was highly efficacious at 3 days across all concentrations (**Fig. 1A**), reflecting the hypersensitivity of these cells to EGFR inhibition. Cytoplasmic vacuoles indicative of autophagy were also observed, consistent with previous reports (28). By day 10, there was marked osimertinib-induced cytotoxicity, with survival reduced to approximately 2–3% of the initial population. At concentrations below 3 nmol/L, a fraction of cells resumed proliferation, whereas at higher concentrations durable proliferative arrest was maintained for at least 3 weeks (Supplementary Fig. S1A). These findings indicate that a single dose of osimertinib induces sustained proliferative arrest in the surviving population, consistent with therapy-induced senescence.

**Figure 1.**
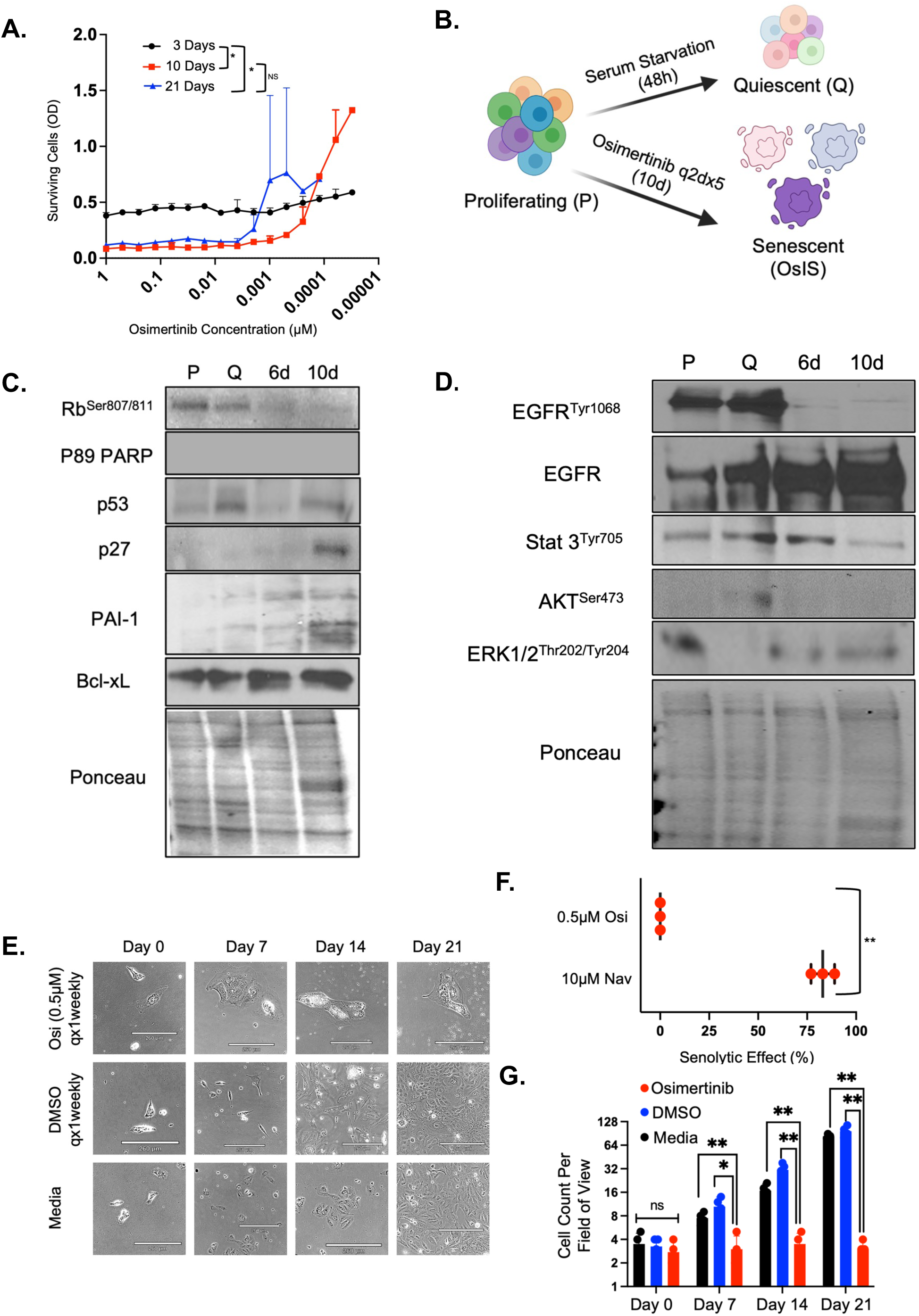
Prolonged Osimertinib Induces Senescence in EGFR+ HCC827 NSCLC. **A.** Cells were treated with 1 μmol/L osimertinib and diluted two-fold across 18 dose levels. Viability was determined at 3, 10, and 21 days using the SRB assay (mean ± SD). Raw SRB absorbances (ODs) that reflect surviving cells, are shown on the Y-axis. **B.** Schema to induce senescence and quiescence. **C.** Immunoblot analysis at 6d and 10d to monitor the timeline of senescence induction. Ponceau confirms equal loading. **D.** Pharmacodynamic evaluation of EGFR pathway components indicating osimertinib-induced suppression of siganling. Ponceau confirms equal loading. **E.** Representative phase-contrast images of OsIS cells exposed to weekly 0.5 μmol/L osimertinib, DMSO vehicle, or media alone. Scale bar corresponds to 260 μm. **F.** Viability of OsIS cells following 72-hour treatment with 0.5 μmol/L osimertinib or 10 μmol/L navitoclax. Individual data points are indicated, together with mean ± SD. **G.** Cell count per field of view of OsIS cells in the presence and absence of osimertinib (mean ± SD) over 21 days. Statistical significance was assessed using paired *t*-test or two-way ANOVA with Tukey’s post-hoc test. *, *P* < 0.05; **, *P* < 0.01; ns, not significant.

To simulate the daily osimertinib dosing that patients receive, repeated daily exposure was evaluated but found to be cytotoxic even at low concentrations (not shown). In contrast, dosing every 48 hours for 10 days killed all but 2 percent of cells, which were non-proliferative. Thus, senescent cells were generated for all experiments by treating HCC827 with 0.5 µmol/L osimertinib for 2 days followed by 0.2 µmol/L every 48 hours until day 10 (**Fig. 1B**). Previous data from our lab and others indicate that ten days of drug exposure is required to maximize cytotoxicity and stabilize senescence (29, 30). Quiescent cells were generated by 48h serum starvation as an alternative proliferative arrest state.

The senescence Associated Beta galactosidase (SA-β-gal) assay (31) was not reliable in HCC827 cells, so senescence was assessed by immunoblot analysis for established markers, collecting lysates at 6d and 10d (**Fig. 1C**). Senescence was confirmed by loss of proliferation (dephosphorylated pRb), absence of apoptosis (PARP), activation of tumor suppressors (TP53 and p27), increased secretion (PAI-1) and increased expression of Bcl-xL (32–34). Henceforth, osimertinib treated cells were termed **Os**imertinib-**I**nduced **S**enescence (OsIS) cells.

The pharmacodynamic effect of osimertinib was confirmed by dephosphorylation of EGFR and its downstream effectors STAT and AKT **(Fig. 1D)**. Consistent with prior reports, ERK1/2 was not fully suppressed by osimertinib in these cells (20). HCC827 cells harbor significant focal EGFR amplification (35) and total EGFR expression was increased in OsIS despite dephosphorylation, suggesting that elevated EGFR copy number is associated with reduced sensitivity to osimertinib-induced cytotoxicity.

Senolytic agents selectively eliminate senescent cells (33). To evaluate whether osimertinib exhibits senolytic activity, OsIS cells were treated weekly with media, DMSO + media or 0.5 µmol/L osimertinib + media and monitored for 21 days (**Fig. 1E**). Osimertinib did not induce cell death in OsIS cells, although autophagic vacuoles were evident indicating a pharmacodynamic effect. In contrast, navitoclax, a validated senolytic (36), killed approximately 80% of OsIS cells (**Fig. 1F** and Supplementary Fig. 1B). In the absence of drug, OsIS cells resumed proliferation by day 7 in both the media-only and DMSO conditions (**Fig. 1E and 1G**), indicating that continued osimertinib is required to maintain the senescent state. Together, these findings demonstrate that osimertinib lacks senolytic activity and instead enforces drug-dependent senescence in HCC827. Similar results were observed in other EGFR+ cell lines and will be reported in a separate manuscript.

### Recurrent Cycles of Senescence Induction and Escape Cause Osimertinib Resistance with Increased Migratory and Invasive Features

To test whether recurrent senescence escape drives acquired resistance, HCC827 cells were subjected to four successive rounds of OsIS with intervening drug withdrawal to enable proliferative recovery (**Fig. 2A**). This generated four independently derived populations (C1–C4) over six to seven months with a cumulative osimertinib exposure of 40 days (10 days per cycle). Senescence-escape populations retained sensitivity to osimertinib until the third selection when cytotoxicity became attenuated (not shown), necessitating higher concentrations to be used in subsequent rounds of selection. Once established, cell lines were maintained without osimertinib. All senescence-escape populations had prolonged doubling time and increased cell area compared to HCC827 (Supplementary Table S2A), indicating phenotypic divergence. Drug sensitivity profiling revealed marked resistance to osimertinib and the 2^nd^ generation EGFR-inhibitor afatinib in all senescence-escape populations **(Fig. 2B and C).** AUC and Emax from dose-response plots are summarized in Supplementary Table. S2B. Overall, response patterns were similar for both inhibitors; although C1 was completely resistant to osimertinib but had weak sensitivity to afatinib, (approximately 20% suppression of proliferation). Like osimertinib, afatinib did not have senolytic activity in OsIS cells (denoted by X in **Fig. 2C**).

**Figure 2.**
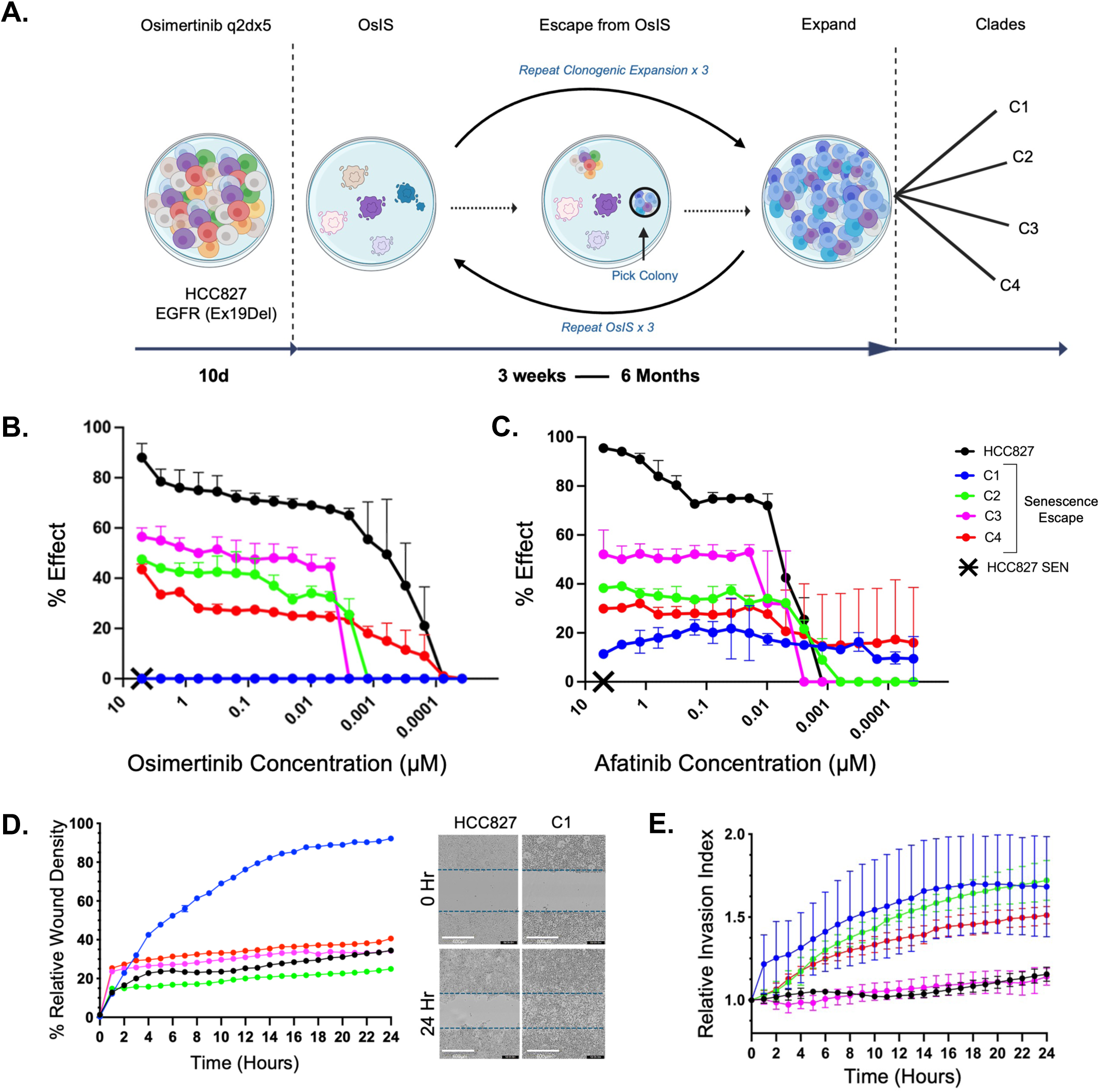
Cells that Escape from Recurrent Senescence Induction are Resistant to Osimertinib. **A.** Schema to generate resistant cell lines via recurrent cycles of senescence induction and escape. **B.** Dose response curves for osimertinib. Cells were treated with 5 μmol/L of drug, diluted two-fold across 18 dose levels and viability assessed after 10 days. Data represents mean ± SD. X denotes the effect of 5 μmol/L osimertinib on OsIS cells. **C.** Dose response curves for afatinib. Cells were treated with 5 μmol/L of drug, diluted two-fold across 18 dose levels and viability assessed after 10 days. Data represents mean ± SD. X denotes the effect of 5 μmol/L afatinib on OsIS cells. **D.** Wound-healing density plot summarizing migratory features of cells (mean ± SD). Representative phase-images of HCC827 and C1 cells at t=0 and 24 hours indicating increased migration of C1. Dashed lines demarcate the position of the wound. Bar represents 600 µm. **E.** Plot quantifying the invasive index of cells under 20% serum gradient (mean ± SD). Data were analyzed according to the manufacturer’s protocol. Two-way ANOVA with Dunnett’s indicate statistically significant difference relative to HCC827 \*\*\*\**P*<0.0001.

Since clinical resistance is closely associated with metastasis, we evaluated the migratory and invasive capacity of senescence-escaped populations. Only C1 showed significantly enhanced wound closure compared with parental cells (**Fig. 2D**), whereas C2, C3, and C4 did not exhibit increased migration (Supplementary Fig. S2). The highest invasion in response to a 20% serum gradient was also observed for C1, followed by intermediate levels for C2 and C4 and minimal invasion for HCC827 and C3 lines (**Fig. 2E**). Thus, the most resistant line, C1 acquires a metastatic-like phenotype characterized by increased migratory and invasive features.

### Recurrent Senescence Escape Causes Chromosomal Instability and Genetic Heterogeneity

To characterize genomic alterations associated with osimertinib resistance, we performed WES and reconstructed a single nucleotide variant SNV-based phylogeny (**Fig. 3A**). All senescence-escape lines shared ancestry with parental HCC827 cells but segregated into distinct evolutionary lineages. C2 was most similar to the parental genotype, followed by C4, whereas C1 and C3 clustered on a divergent branch. The most osimertinib-resistant line C1 had the greatest genetic divergence. Hereafter, we refer to these senescence-escape lineages as clades.

**Figure 3.**
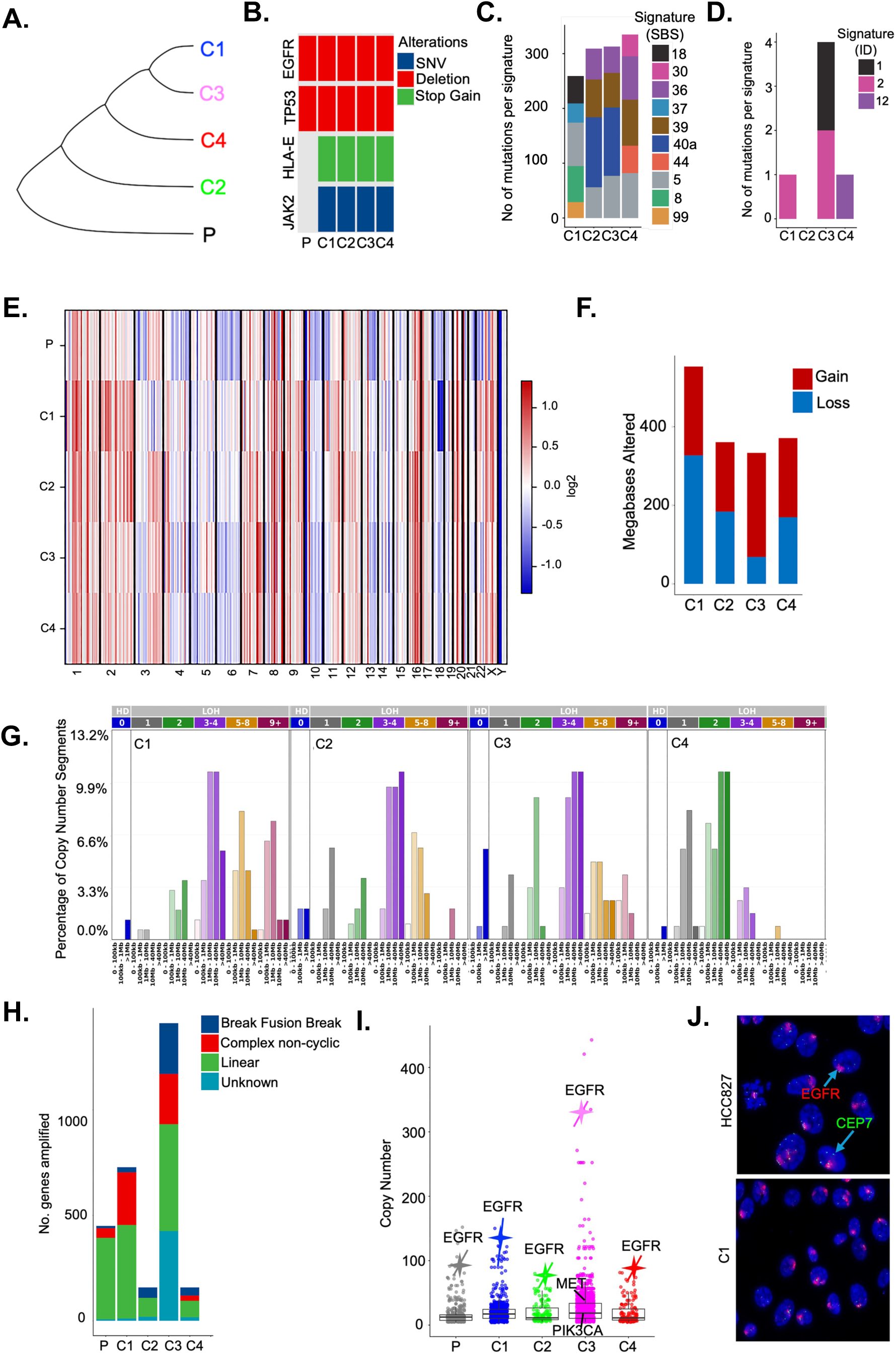
Recurrent Senescence Escape Causes Genomic Heterogeneity via Chromosomal Instability. **A.** Phylogenetic reconstruction of HCC827 and senescence-escape clades based on Single-Nucleotide Variants (SNVs) from WES. **B.** Focused oncoplot illustrating representative somatic variants. Type of SNV is indicated. **C.** Histogram plot showing single-base substitution (SBS) mutational signatures normalized to HCC827. **D.** Histogram plot showing indel (ID) mutational signatures normalized to HCC827. **E.** Heatmap showing genome-wide copy number variation analysis of cell lines. The intensity of color reflects the magnitude of change. Red indicates copy number gain and blue indicates loss. **F.** Histogram plot showing the size of megabases altered by somatic CNVs in each clade. Data is normalized to HCC827. **G.** Histogram plot showing allele-specific copy number distributions indicating large scale LOH across clades, particularly in C1. **H.** WGS and amplicon architecture classification of amplicon structures across cell lines. **I.** Scatter plot of genes amplified across all cell lines indicating different distribution. Boxplot indicates median ± IQR. *EGFR* copy-number are indicated for each cell line. Copy number amplification for *MET* and *PIK3CA* are indicated for C3. **J.** Representative FISH images of HCC827 and C1 cells showing extensive *EGFR* amplification (red) visible as HSR’s. Chromosome 7 centromere, CEP7 is green. Magnification, 100×.

A focused oncoplot of representative somatic variants is shown in **Fig. 3B**. *De novo* SNVs or indels in *EGFR* were not observed, suggesting that resistance to osimertinib was not driven by on-target mutations. Neither HCC827 nor the clades expressed wild-type *EGFR,* eliminating this as a potential resistance mechanism. Similarly, *de novo* mutations were not observed in *TP53*. Novel SNVs with high functional priority were detected in all clades, consistent with ongoing mutational diversification during repeated cycles of OsIS and escape. HCC827 cells harbor *HLA-C* missense and *HLA-B* synonymous variants, and all clades acquired a *de novo* stop-gain mutation in *HLA-E*, suggesting an attenuated immunogenic effect. A variant in *JAK2*, a downstream effector of EGFR signaling, was also present in all clades, suggesting acquisition early in selection. All clades had increased tumor mutational burden (TMB) compared to the parent line (Supplementary Fig. S3A). SNVs common to all clades and unique to each are summarized in Supplementary Tables S2. Sequencing depth and SNV and indel frequencies are summarized in Supplementary Tables S3A-C.

Mutational signature analysis (26) revealed complex patterns of single nucleotide substitutions (SBS) (**Fig. 3C**, Supplementary Fig. S3B & Table S3D). All clades had an age-associated clock-like signature SBS5, consistent with background mutational accumulation. The most resistant clade, C1 had a distinct mutational signature profile enriched for SBS18, SBS99, SBS37 and SBS8 indicating oxidative DNA damage and melphalan mutagenesis. C4 had signatures reflecting defective DNA mismatch repair (SBS44) and base excision repair due to MUTYH deficiency and NTHL1 inactivation (SBS36 and 30, respectively). C2 and C3 had the same mutational signatures, enriched for defective base excision repair (SBS36). Insertion-deletion (indel) signature analysis identified replication slippage signatures, ID1 and ID2, and an unknown signature **(Fig. 3D**, Supplementary Fig. S3C & Table S3D), although rare relative to SBS burden, indicating that point mutagenesis was dominant. The relative enrichment of these signatures across clades is consistent with ongoing mutagenesis during repeated cycles of senescence induction and escape.

#### Copy Number Variation Analysis

Genome-wide copy number analysis revealed a highly variable landscape with significant divergence (**Fig. 3E**). Quantification of copy-number alterations megabases of altered genome among clades (**Fig. 3F**). C1 had the highest copy-number alterations driven predominantly by loss, while C3 displayed the lowest overall change, though it had the highest copy-number gains. Copy number variations in genes identified from genomic analysis of ctDNA derived from resistant patients (7) were not detected. This was also confirmed at the protein level by immunoblotting (Supplementary Fig. S3D). To characterize LOH across the genomic landscape, we analyzed allele specific genomic features. As shown in **Fig. 3G**, copy number alteration segment distributions were dominated by large-scale LOH events particularly for C1, consistent with a high level of chromosomal instability.

To define the structural organization of copy-number alterations, we used AmpliconArchitect to reconstruct amplifications and highly rearranged genomic regions from WGS (37). Distinct amplification architectures were observed across parental and resistant clades **(Fig. 3H)**. The most resistant clade, C1 exhibited an architecture dominated by linear amplification with complex non-cyclic structures. C3 had the highest number of amplified genes and was significantly enriched for breakage-fusion-bridge and complex non-cyclic amplicons. In contrast, C2 and C4 were largely characterized by linear amplification topologies. Breakage-fusion-bridge (BFB) cycles, indicating chromosomal instability were increased in all clades.

EGFR amplification was highest in C3 rather than the most resistant clade C1 (**Fig. 3I**). Concordance between WES and WGS copy-number estimates was limited to *EGFR* and *CDK4* amplifications only (Supplementary Table S3E). Therefore, neither increased global amplification or *EGFR* copy number correlated with resistance. Low-level *MET* amplification was only detected in C3 by AmpliconArchitect (**Fig. 3I** & Supplementary Table S3E) and did not translate into increased MET protein expression (Supplementary Fig. S3D). Notably, MET expression was significantly reduced in the most resistant clade C1, indicating that amplification does not correlate with resistance. FISH analysis of *EGFR* and CEP7 also demonstrated extensive *EGFR* amplification evident as homogeneously staining regions (35). Aneuploidy was evident across all lines; however, copy number could not be determined since the *EGFR* signal exceeded the threshold for accurate quantitation (**Fig. 3J**). Together, these data indicate that osimertinib resistance via recurrent senescent escape emerges without *de novo* EGFR mutations. Chromosomal instability was the most prevalent genomic feature that correlated with resistance, consistent with the majority of patients upon relapse.

### Resistant Cells Retains Efficacy to Platinum-Based Chemotherapy

We next assessed sensitivity to standard-of-care platinum–pemetrexed chemotherapy (4). Cisplatin had comparable efficacy across all clades **(Fig. 4A)**, whereas pemetrexed sensitivity was variable, with C1 exhibiting marked resistance **(Fig. 4B)**. Immunoblotting for thymidylate synthase, predictive of pemetrexed sensitivity (38) revealed no difference in expression across clades **(Fig. 4C)**. AUC and Emax from dose-response plots are summarized in Supplementary Table S4A. Importantly, cisplatin killed approximately 70% of OsIS cells, whereas pemetrexed had no effect, confirming our prior observations that cisplatin is a senolytic (29, 39).

**Figure 4.**
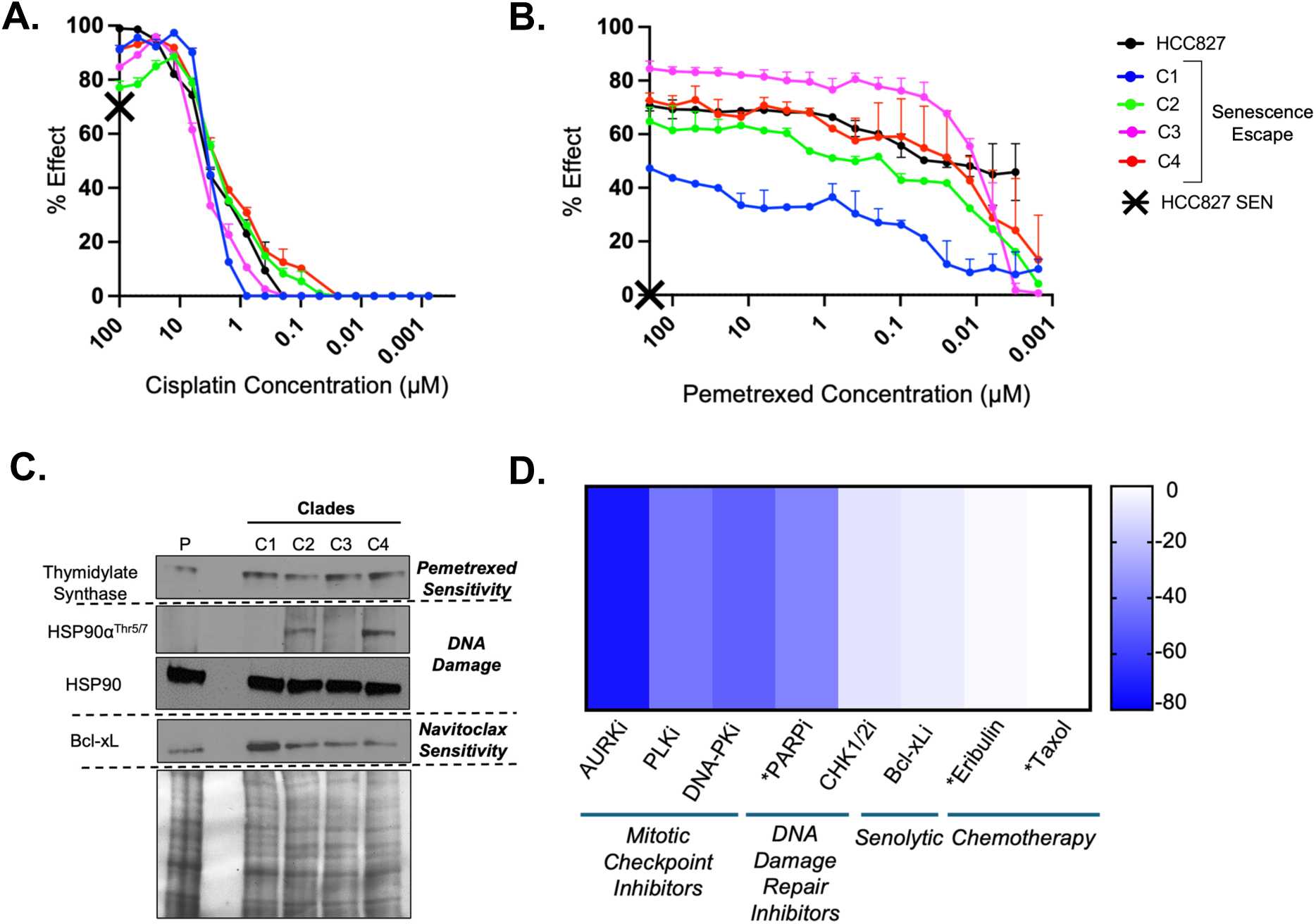
Osimertinib-Resistant Cells Retain Sensitivity to Chemotherapy. **A.** Dose response curves for cisplatin. Cells were treated with 100 μmol/L of drug diluted two-fold across 18 dose levels, and viability assessed after 10 days. Data represents mean ± SD. X denotes approximately 70% effect of 100 μmol/L cisplatin on OsIS cells, indicating that cisplatin is a senolytic. Curves were not statistically significantly different from HCC827 (two-way ANOVA with Tukey’s post-hoc test), except C3 *, *P* < 0.05. **B.** Dose response curves for pemetrexed. Cells were treated with 200 μmol/L of drug diluted two-fold across 18 dose levels, and viability assessed after 10 days. Data represents mean ± SD. X denotes the effect of 200 μmol/L pemetrexed on OsIS cells. Curves were statistically significantly different from HCC827 (two-way ANOVA with Tukey’s post-hoc test, ****, *P* < 0.0001), except C4, ns. **C.** Immunoblot analysis of proteins potentially predictive of drug response. Ponceau staining confirms equal protein loading. **D.** Heatmap showing relative changes in drug response of C1 compared to HCC827. Area under the curve (AUC) was calculated from dose-response curves and relative change in C1 calculated. White indicates comparable sensitivity while blue indicates resistance (decreased AUC in C1). * indicates FDA-approved drugs.

Given the extensive genomic instability of C1, we investigated its therapeutic vulnerabilities to mitotic checkpoint inhibitors (barasertib and volasertib), DNA damage response inhibitors (olaparib, AZD7648, and AZD7762), the Bcl-xL inhibitor navitoclax and tubulin-targeting chemotherapy (Taxol and eribulin). Relative AUC analysis showed significant resistance to mitotic checkpoint inhibitors and moderate resistance to DNA repair inhibitors (**Fig. 4D** and Supplementary Fig. S4A). Notably, C1 was resistant to the DNA-PK inhibitor AZD7648 despite pronounced genomic instability; however, phosphorylation of HSP90 at the DNA-PK-dependent site was unchanged (**Fig. 4C**), indicating that resistance was independent of DNA-PK pathway activation.

In contrast, sensitivity to tubulin-targeting drugs (Taxol and eribulin) was retained, consistent with the clinical observation that taxanes have activity in osimertinib-refractory patients (40). Navitoclax, which has been clinically evaluated in combination with osimertinib (41), also retained sensitivity in C1 cells, which coincides with the increased Bcl-xL expression, as shown in **Fig. 4C**. Collectively, these results indicate that osimertinib resistance associated with senescence escape is not overcome by targeting mitotic checkpoints or DNA repair pathways, but is susceptible to platinum- and tubulin-targeting chemotherapy.

## Discussion

Acquired resistance to osimertinib is nearly universal in EGFR+ lung cancer, yet only a minority of cases are explained by on-target EGFR mutations. By modeling resistance through recurrent cycles of osimertinib-induced senescence and escape rather than continuous dose escalation, we recapitulate a clinically relevant evolutionary trajectory characterized by prolonged dormancy followed by relapse. A critical finding is that continuous osimertinib maintains senescence without eliminating established senescent cells, whereas drug withdrawal permits proliferative escape. This has direct clinical relevance, as effective osimertinib exposure fluctuates due to heterogeneous intratumoral penetration, variable systemic distribution, and treatment interruptions related to toxicity, adherence, or access. Collectively, these results establish senescence not as a terminal tumor-suppressive state, but as a drug-tolerant survival program that acts as an evolutionary bottleneck from which resistant populations emerge.

Notably, our group previously reported that cisplatin has senolytic activity in taxol- or cisplatin-induced senescent cancer cells (29, 39). Similarly, we show here that cisplatin kills OsIS cells, and that osimertinib-resistant cells remain responsive to cisplatin. This is consistent with clinical observations and provides mechanistic insight that could explain the superior clinical efficacy of platinum-based combinations in EGFR-mutant disease (6). Although resistant populations respond to taxanes, platinum-pemetrexed regimens have better outcomes suggesting that the ability to eliminate senescent cells may be as important as cytotoxic potency against proliferating tumors.

Resistance driven by recurrent senescence escape was marked by chromosomal instability and genome-scale disorder rather than discrete gene-level alterations; however, despite profound chromosomal instability, resistant clades were refractory to mitotic checkpoint and DNA repair inhibitors, underscoring the clinical reality that genomic disorder does not necessarily confer therapeutic vulnerability. Rather, once tumors adapt to tolerate checkpoint dysfunction and genomic chaos, targeting these pathways is ineffective. This has also been borne out by the fact that molecules targeting these cellular functions have not been FDA-approved due to lack of efficacy and / or toxicity. Together, these data support models in which sustained therapeutic pressure fuels genomic entropy, as shown in lung cancer by TRACERx (42) and chromosomal instability acts a common driver of resistance and progression across malignancies (43).

In addition to the population bottleneck imposed by the small fraction of cells that survive osimertinib treatment, senescence itself may also function as a mutagenic bottleneck. Senescent cells harbor persistent DNA damage and have impaired DNA repair function (44). They also frequently exhibit whole-genome doubling and aneuploidy (45, 46) and as recently documented, acquire *de novo* structural rearrangements such as chromosomal inversions that directly enable senescence escape (47). Although the prevalence and kinetics of these processes remain to be fully defined *in vivo*, these observations position senescence as an active evolutionary state that promotes chromosomal instability and genome diversification under therapeutic pressure.

Ultimately, these data underscore the central role of senescent tumor cells in driving cancer evolution and resistance. Overexpression of Bcl-xL is a reproducible feature of senescent and resistant cancer cells and tumors, suggesting mechanistic relevance; however, clinical and preclinical efforts to target this pathway have been constrained by thrombocytopenia and limited efficacy with navitoclax and related inhibitors (41, 48). Although senolytic strategies have shown limited clinical success in aging-related contexts, cancers that have prolonged responses to targeted therapy represent clinically tractable opportunities to test senolytic adjuvants and emerging clinical efforts support the feasibility of this approach (49). In summary, our findings identify senescence escape as a key mechanism of osimertinib resistance and suggest that durable disease control will require not only sustained oncogenic pathway inhibition, but early elimination of dormant tumor reservoirs before genomic instability and relapse develop.

## Authors’ Contributions

N. Jamil: Data curation, analysis, visualization, validation, investigation, methodology, writing original draft, writing review and editing. Q. Hooker: Data curation, formal analysis, investigation, visualization, methodology. N. Cornejal: investigation, data curation and methodology. H.D. Hosgood: supervision. H. McDaid: Conceptualization, resources, formal analysis, supervision, funding acquisition, methodology, writing original draft, project administration, writing review and editing.

## Acknowledgments

The authors thank Sumaiya Jessi (Univ. Buffalo), and the Einstein Cytogenetic core facility. This work was supported in part by the US Department of Defense Concept award W81XWH-20-1-0499 (HMD), National Cancer Institute 1R01CA279952 (HMD) and the Breast Cancer Research Foundation (HMD). N.Jamil was supported by the National Center for Advancing Translational Sciences predoctoral program in Clinical Investigation, T32TR004537 (HDH). Cores services were supported by the Montefiore Einstein Comprehensive Cancer Center funding, 5P30CA013330-53. The content is solely the responsibility of the authors and does not necessarily represent the official views of the National Institutes of Health. AI was used to assist with editing.

## Note

Supplementary data for this article are available at Cancer Research Online (http://cancerres.aacrjournals.org/).

## REFERENCES

1. Soria JC, Ohe Y, Vansteenkiste J, Reungwetwattana T, Chewaskulyong B, Lee KH, et al. Osimertinib in Untreated EGFR-Mutated Advanced Non-Small-Cell Lung Cancer. N Engl J Med. 2018;378(2):113–25.

2. Ramalingam SS, Vansteenkiste J, Planchard D, Cho BC, Gray JE, Ohe Y, et al. Overall Survival with Osimertinib in Untreated, EGFR-Mutated Advanced NSCLC. New England Journal of Medicine. 2020;382(1):41–50.

3. Planchard D, Janne PA, Cheng Y, Yang JC, Yanagitani N, Kim SW, et al. Osimertinib with or without Chemotherapy in EGFR-Mutated Advanced NSCLC. N Engl J Med. 2023;389(21):1935–48.

4. Janne PA, Planchard D, Kobayashi K, Yang JC, Liu Y, Valdiviezo N, et al. Survival with Osimertinib plus Chemotherapy in EGFR-Mutated Advanced NSCLC. N Engl J Med. 2025.

5. Patil T, Gao D, Watson A, Sakamoto M, Nie Y, Gibson A, et al. The eaicacy of continuing osimertinib with platinum pemetrexed chemotherapy upon progression in patients with metastatic non-small cell lung cancer harboring sensitizing EGFR mutations. Lung Cancer. 2025;199:108040.

6. Peled N, Tufman A, Sequist LV, Pasello G, Wang Q, Antonuzzo L, et al. COMPEL: osimertinib plus platinum-based chemotherapy in patients with EGFR-mutated advanced NSCLC and progression on first-line osimertinib. ESMO Open. 2025;10(10):105807.

7. Chmielecki J, Gray JE, Cheng Y, Ohe Y, Imamura F, Cho BC, et al. Candidate mechanisms of acquired resistance to first-line osimertinib in EGFR-mutated advanced non-small cell lung cancer. Nat Commun. 2023;14(1):1070.

8. Gray JE, Markovets A, Reungwetwattana T, Majem M, Nogami N, Peled N, et al. Longitudinal Analyses of Circulating Tumor DNA for the Detection of EGFR Mutation-Positive Advanced NSCLC Progression During Treatment: Data From FLAURA and AURA3. J Thorac Oncol. 2024;19(11):1525–38.

9. Zhan J, Xue J, Wu L, Zhang Z, Wang Q, Ma Y, et al. HS-10375, a selective EGFR C797S tyrosine kinase inhibitor, in advanced non-small cell lung cancer. J Transl Med. 2025;23(1):628.

10. Chmielecki J, Mok T, Wu YL, Han JY, Ahn MJ, Ramalingam SS, et al. Analysis of acquired resistance mechanisms to osimertinib in patients with EGFR-mutated advanced non-small cell lung cancer from the AURA3 trial. Nat Commun. 2023;14(1):1071.

11. Sequist LV, Han JY, Ahn MJ, Cho BC, Yu H, Kim SW, et al. Osimertinib plus savolitinib in patients with EGFR mutation-positive, MET-amplified, non-small-cell lung cancer after progression on EGFR tyrosine kinase inhibitors: interim results from a multicentre, open-label, phase 1b study. Lancet Oncol. 2020;21(3):373–86.

12. de Marinis F, Kim TM, Bonanno L, Cheng S, Kim SW, Tiseo M, et al. Savolitinib plus osimertinib in epidermal growth factor receptor (EGFR)-mutated advanced non-small cell lung cancer with MET overexpression and/or amplification following disease progression on osimertinib: primary results from the phase II SAVANNAH study. Ann Oncol. 2025;36(8):920–33.

13. McGranahan N, Swanton C. Clonal Heterogeneity and Tumor Evolution: Past, Present, and the Future. Cell. 2017;168(4):613–28.

14. Tsuboi M, Herbst RS, John T, Kato T, Majem M, Grohe C, et al. Overall Survival with Osimertinib in Resected EGFR-Mutated NSCLC. N Engl J Med. 2023;389(2):137–47.

15. Hanahan D. Hallmarks of Cancer: New Dimensions. Cancer Discov. 2022;12(1):31–46.

16. Saleh T, Alhesa A, Al-Balas M, Abuelaish O, Mansour A, Awad H, et al. Expression of therapy-induced senescence markers in breast cancer samples upon incomplete response to neoadjuvant chemotherapy. Biosci Rep. 2021;41(5).

17. te Poele RH, Okorokov AL, Jardine L, Cummings J, Joel SP. DNA Damage Is Able to Induce Senescence in Tumor Cells in Vitro and in Vivo1. Cancer Research. 2002;62(6):1876–83.

18. Wang M, Morsbach F, Sander D, Gheorghiu L, Nanda A, Benes C, et al. EGF receptor inhibition radiosensitizes NSCLC cells by inducing senescence in cells sustaining DNA double-strand breaks. Cancer Res. 2011;71(19):6261–9.

19. Gerber PA, Buhren BA, Schrumpf H, Hevezi P, Bölke E, Sohn D, et al. Mechanisms of skin aging induced by EGFR inhibitors. Support Care Cancer. 2016;24(10):4241–8.

20. Kurppa KJ, Liu Y, To C, Zhang T, Fan M, Vajdi A, et al. Treatment-Induced Tumor Dormancy through YAP-Mediated Transcriptional Reprogramming of the Apoptotic Pathway. Cancer Cell. 2020;37(1):104–22 e12.

21. Chao SK, Lin J, Brouwer-Visser J, Smith AB, 3rd, Horwitz SB, McDaid HM. Resistance to discodermolide, a microtubule-stabilizing agent and senescence inducer, is 4E-BP1-dependent. Proc Natl Acad Sci U S A. 2011;108(1):391–6.

22. Milanovic M, Fan DNY, Belenki D, Däbritz JHM, Zhao Z, Yu Y, et al. Senescence-associated reprogramming promotes cancer stemness. Nature. 2018;553(7686):96–100.

23. Shien K, Toyooka S, Yamamoto H, Soh J, Jida M, Thu KL, et al. Acquired resistance to EGFR inhibitors is associated with a manifestation of stem cell-like properties in cancer cells. Cancer Res. 2013;73(10):3051–61.

24. Starble RM, Sun EG, Gbyli R, Radda J, Lu J, Jensen TB, et al. Epigenetic priming promotes tyrosine kinase inhibitor resistance and oncogene amplification. Nature Structural & Molecular Biology. 2025.

25. Vichai V, Kirtikara K. Sulforhodamine B colorimetric assay for cytotoxicity screening. Nature Protocols. 2006;1(3):1112–6.

26. Díaz-Gay M, Vangara R, Barnes M, Wang X, Islam SMA, Vermes I, et al. Assigning mutational signatures to individual samples and individual somatic mutations with SigProfilerAssignment. Bioinformatics. 2023;39(12).

27. Shen R, Seshan VE. FACETS: allele-specific copy number and clonal heterogeneity analysis tool for high-throughput DNA sequencing. Nucleic Acids Res. 2016;44(16):e131.

28. Wei Y, Zou Z, Becker N, Anderson M, Sumpter R, Xiao G, et al. EGFR-Mediated Beclin 1 Phosphorylation in Autophagy Suppression, Tumor Progression, and Tumor Chemoresistance. Cell. 2013;154(6):1269–84.

29. Samaraweera L, Adomako A, Rodriguez-Gabin A, McDaid HM. A Novel Indication for Panobinostat as a Senolytic Drug in NSCLC and HNSCC. Sci Rep. 2017;7(1):1900.

30. Young AR, Narita M, Ferreira M, Kirschner K, Sadaie M, Darot JF, et al. Autophagy mediates the mitotic senescence transition. Genes Dev. 2009;23(7):798–803.

31. Debacq-Chainiaux F, Erusalimsky JD, Campisi J, Toussaint O. Protocols to detect senescence-associated beta-galactosidase (SA-betagal) activity, a biomarker of senescent cells in culture and in vivo. Nat Protoc. 2009;4(12):1798–806.

32. Kirkland JL, Tchkonia T. Cellular Senescence: A Translational Perspective. EBioMedicine. 2017;21:21–8.

33. Zhu Y, Tchkonia T, Pirtskhalava T, Gower AC, Ding H, Giorgadze N, et al. The Achilles’ heel of senescent cells: from transcriptome to senolytic drugs. Aging Cell. 2015;14(4):644–58.

34. Kortlever RM, Higgins PJ, Bernards R. Plasminogen activator inhibitor-1 is a critical downstream target of p53 in the induction of replicative senescence. Nat Cell Biol. 2006;8(8):877–84.

35. Raeisi Dehkordi S, Wong IT-L, Ni J, Luebeck J, Zhu K, Prasad G, et al. Breakage fusion bridge cycles drive high oncogene number with moderate intratumoural heterogeneity. Nature Communications. 2025;16(1):1497.

36. Zhu Y, Tchkonia T, Fuhrmann-Stroissnigg H, Dai HM, Ling YY, Stout MB, et al. Identification of a novel senolytic agent, navitoclax, targeting the Bcl-2 family of anti-apoptotic factors. Aging Cell. 2016;15(3):428–35.

37. Deshpande V, Luebeck J, Nguyen ND, Bakhtiari M, Turner KM, Schwab R, et al. Exploring the landscape of focal amplifications in cancer using AmpliconArchitect. Nat Commun. 2019;10(1):392.

38. Paradiso A, Simone G, Petroni S, Leone B, Vallejo C, Lacava J, et al. Thymidilate synthase and p53 primary tumour expression as predictive factors for advanced colorectal cancer patients. Br J Cancer. 2000;82(3):560–7.

39. Guo B, Rodriguez-Gabin A, Prota AE, Mühlethaler T, Zhang N, Ye K, et al. Structural Refinement of the Tubulin Ligand (+)-Discodermolide to Attenuate Chemotherapy-Mediated Senescence. Molecular Pharmacology. 2020;98(2):156–67.

40. Herbst RS, Prager D, Hermann R, Fehrenbacher L, Johnson BE, Sandler A, et al. TRIBUTE: a phase III trial of erlotinib hydrochloride (OSI-774) combined with carboplatin and paclitaxel chemotherapy in advanced non-small-cell lung cancer. J Clin Oncol. 2005;23(25):5892–9.

41. Bertino EM, Gentzler RD, Cliaord S, Kolesar J, Muzikansky A, Haura EB, et al. Phase IB Study of Osimertinib in Combination with Navitoclax in EGFR-mutant NSCLC Following Resistance to Initial EGFR Therapy (ETCTN 9903). Clin Cancer Res. 2021;27(6):1604–11.

42. Frankell AM, Dietzen M, Al Bakir M, Lim EL, Karasaki T, Ward S, et al. The evolution of lung cancer and impact of subclonal selection in TRACERx. Nature. 2023;616(7957):525–33.

43. Chen X, Agustinus AS, Li J, DiBona M, Bakhoum SF. Chromosomal instability as a driver of cancer progression. Nature Reviews Genetics. 2025;26(1):31–46.

44. Rodier F, Muñoz DP, Teachenor R, Chu V, Le O, Bhaumik D, et al. DNA-SCARS: distinct nuclear structures that sustain damage-induced senescence growth arrest and inflammatory cytokine secretion. J Cell Sci. 2011;124(Pt 1):68–81.

45. Dewhurst SM, McGranahan N, Burrell RA, Rowan AJ, Grönroos E, Endesfelder D, et al. Tolerance of whole-genome doubling propagates chromosomal instability and accelerates cancer genome evolution. Cancer Discov. 2014;4(2):175–85.

46. Zhang L, De Cecco M, Lee M, Hao X, Maslov AY, Montagna C, et al. Analysis of Somatic Mutations in Senescent Cells Using Single-Cell Whole-Genome Sequencing. Aging Biol. 2023;1(1).

47. Zampetidis CP, Galanos P, Angelopoulou A, Zhu Y, Polyzou A, Karamitros T, et al. A recurrent chromosomal inversion suaices for driving escape from oncogene-induced senescence via subTAD reorganization. Molecular Cell. 2021;81(23):4907–23.e8.

48. Crespo-Garcia S, Fournier F, Diaz-Marin R, Klier S, Ragusa D, Masaki L, et al. Therapeutic targeting of cellular senescence in diabetic macular edema: preclinical and phase 1 trial results. Nature Medicine. 2024;30(2):443–54.

49. DeMichele A, Clark AS, Shea E, Bayne LJ, Sterner CJ, Rohn K, et al. Targeting dormant tumor cells to prevent recurrent breast cancer: a randomized phase 2 trial. Nature Medicine. 2025;31(10):3464–74.

