## Supplementary Figures for "Recurrent Escape from Osimertinib-Induced Senescence Promotes Genomic Instability Associated with Therapeutic Resistance"

### Slide 1
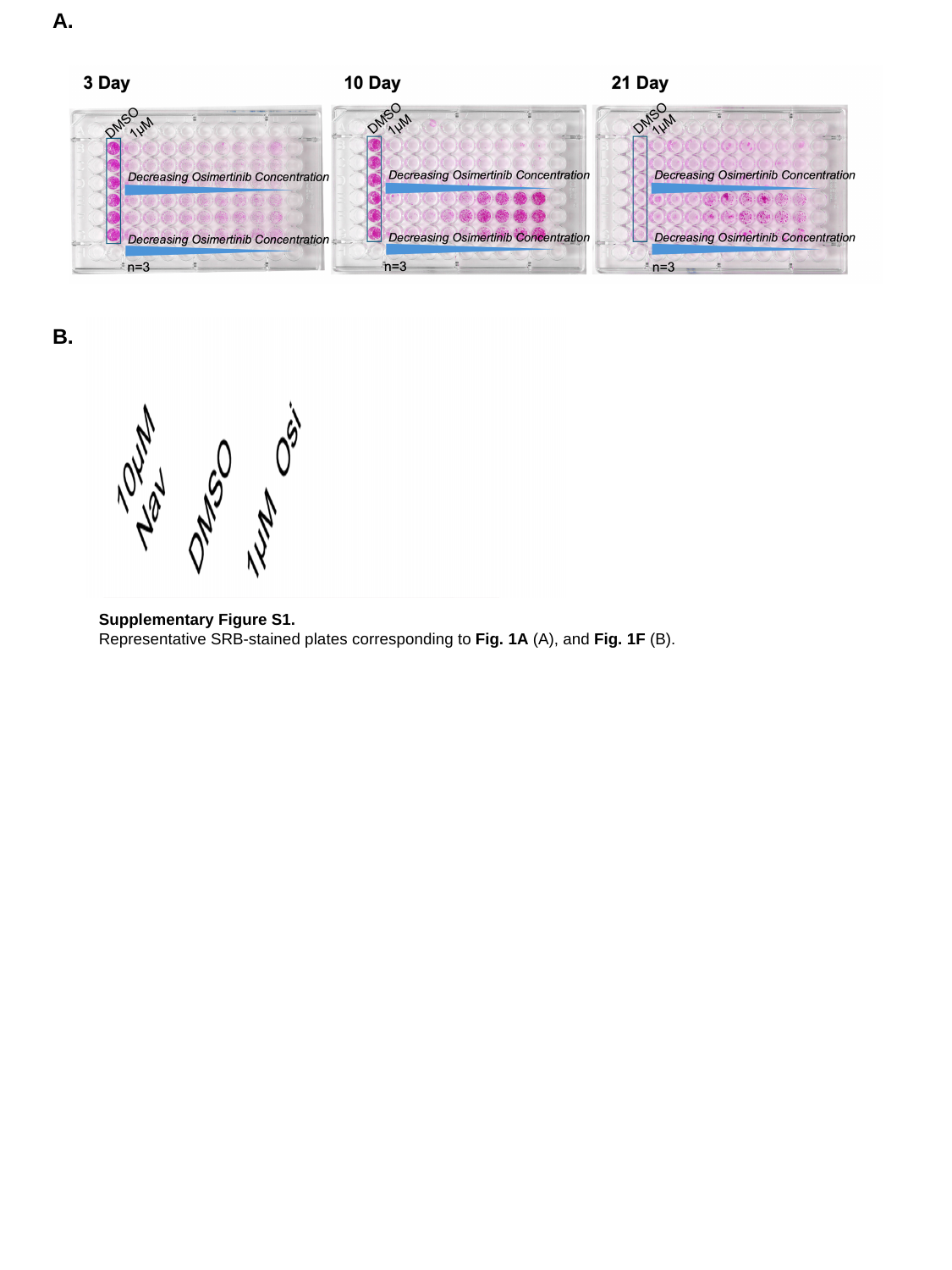

A.
B.
Supplementary Figure S1.
Representative SRB-stained plates corresponding to Fig. 1A (A), and Fig. 1F (B).

### Slide 2
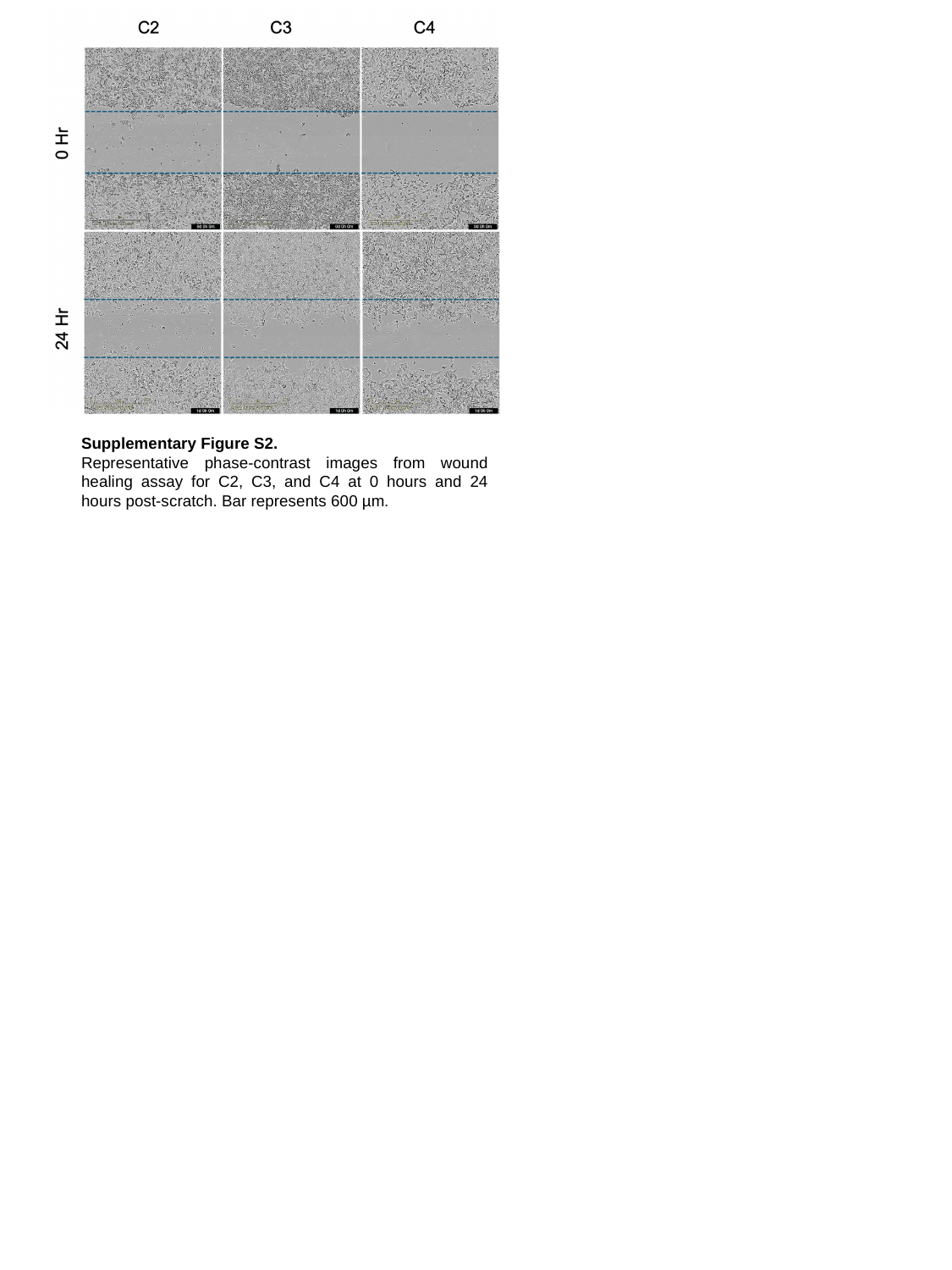

Supplementary Figure S2.
Representative phase-contrast images from wound healing assay for C2, C3, and C4 at 0 hours and 24 hours post-scratch. Bar represents 600 µm.

### Slide 3
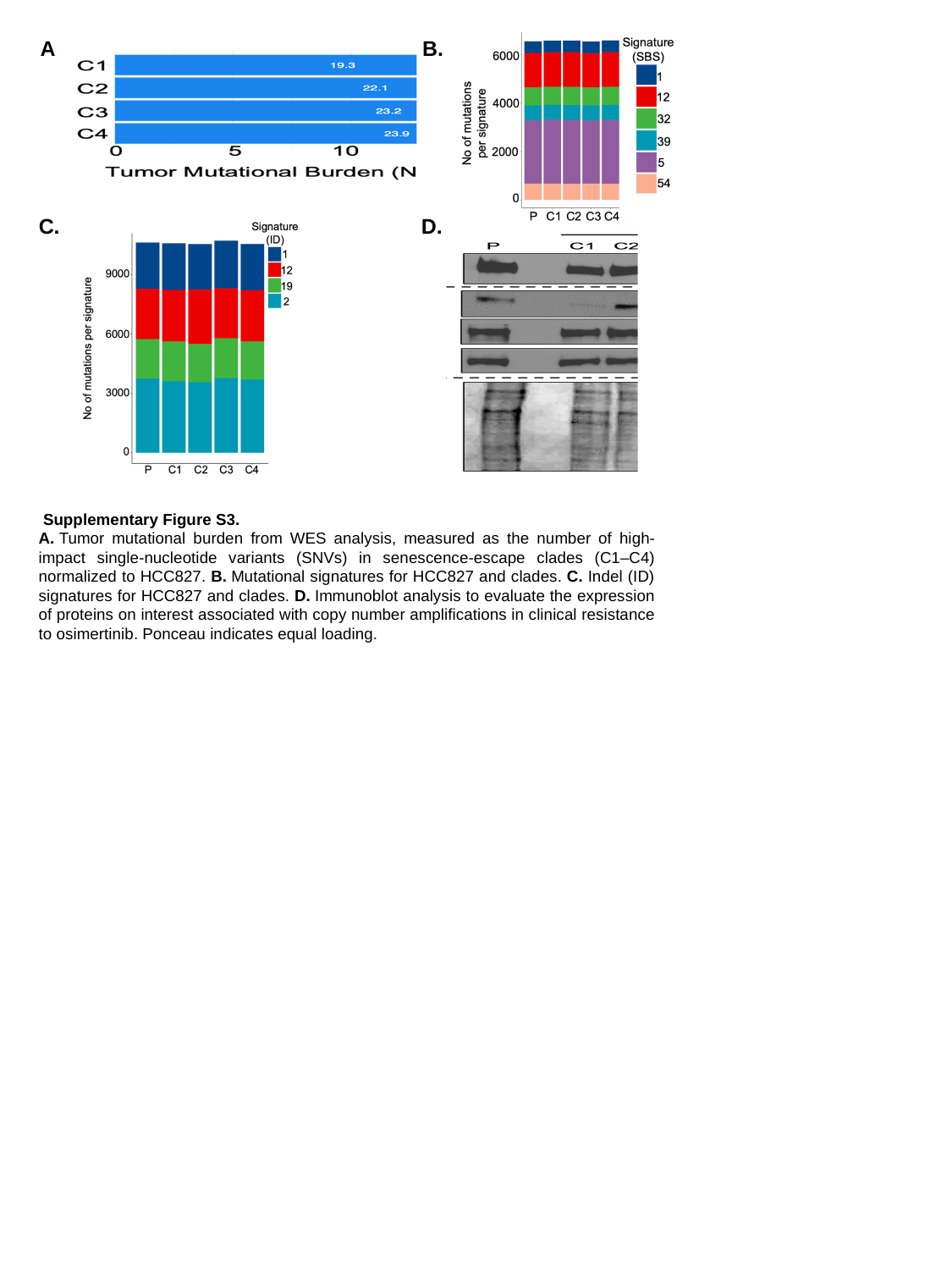

B.
A.
C.
D.
 Supplementary Figure S3.
A. Tumor mutational burden from WES analysis, measured as the number of high-impact single-nucleotide variants (SNVs) in senescence-escape clades (C1–C4) normalized to HCC827. B. Mutational signatures for HCC827 and clades. C. Indel (ID) signatures for HCC827 and clades. D. Immunoblot analysis to evaluate the expression of proteins on interest associated with copy number amplifications in clinical resistance to osimertinib. Ponceau indicates equal loading.

### Slide 4
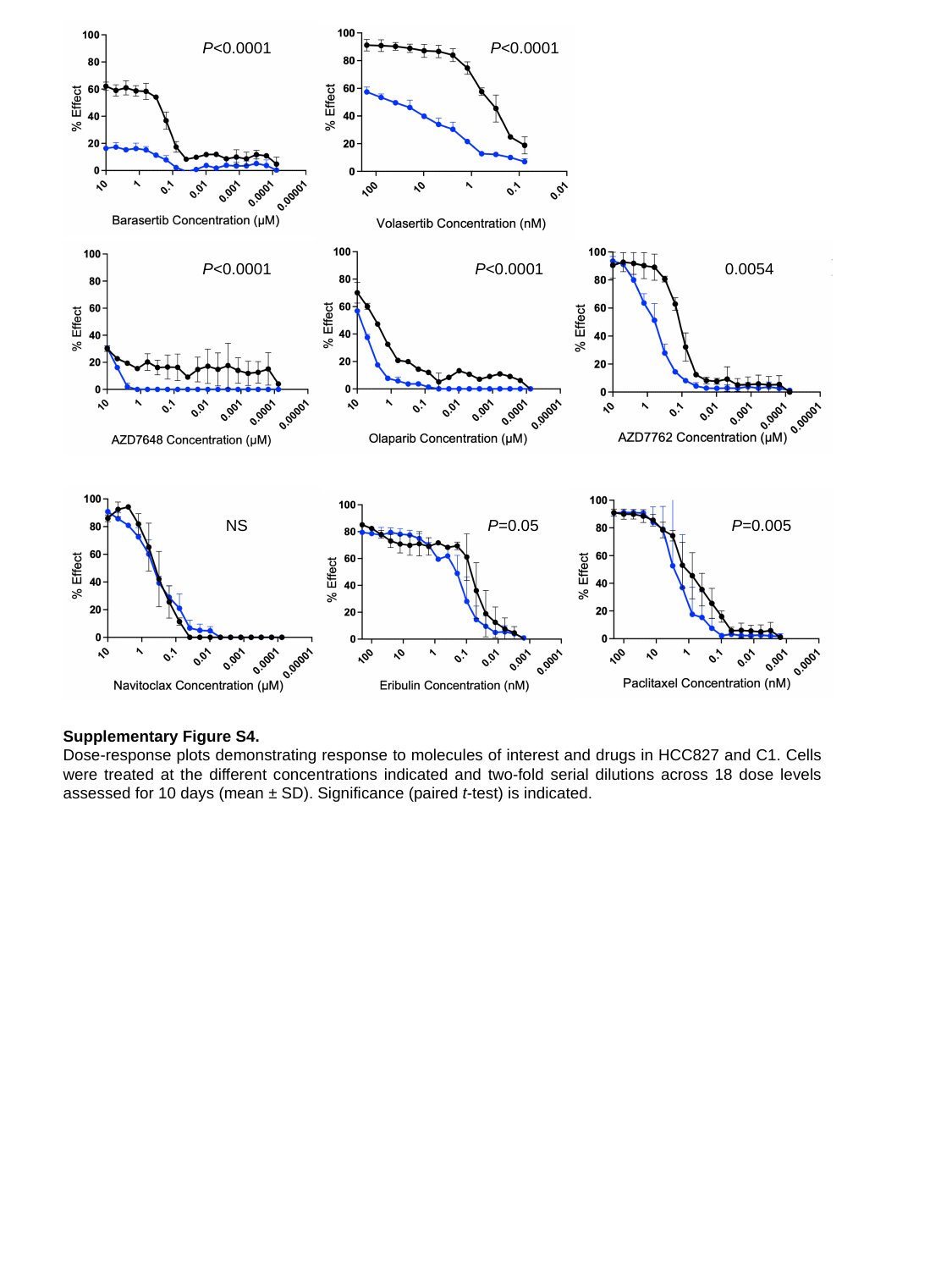

P<0.0001
P<0.0001
P<0.0001
P<0.0001
0.0054
NS
P=0.05
P=0.005
Supplementary Figure S4.
Dose-response plots demonstrating response to molecules of interest and drugs in HCC827 and C1. Cells were treated at the different concentrations indicated and two-fold serial dilutions across 18 dose levels assessed for 10 days (mean ± SD). Significance (paired t-test) is indicated.
